# Cross-seasonal weather effects interact with breeding conditions to impact reproductive success in an alpine songbird

**DOI:** 10.1101/2021.08.06.455393

**Authors:** Devin R. de Zwaan, Anna Drake, Alaine F. Camfield, Elizabeth C. MacDonald, Kathy Martin

## Abstract

1. In alpine habitats, fluctuating early-season weather conditions and short breeding seasons limit reproductive opportunities, such that arriving and breeding earlier or later than the optimum may be particularly costly for migratory species. Given early-season energy limitations, the influence of environmental conditions across the annual cycle on breeding phenology may have pronounced fitness consequences, yet our understanding of cross-seasonal dynamics in alpine breeding organisms is severely limited.
2. For an alpine-breeding, migratory population of horned lark (*Eremophila alpestris*) in northern British Columbia, Canada (54.8°N latitude) we assessed how spatially explicit weather conditions from across the annual cycle influenced clutch initiation date and offspring development. We also addressed how cross-seasonal effects on breeding parameters interact to influence reproductive fitness.
3. With 12 years of intensive breeding data and 3 years of migration data from archival light-level geolocators, we used a sliding window approach to identify critical points during the annual cycle where weather events most influenced breeding phenology and offspring development. Consequences for reproductive success were assessed using nest survival simulations.
4. Average clutch initiation varied up to 11 days among years but did not advance from 2003 to 2019. Colder temperatures with greater precipitation at wintering habitats, as well as colder temperatures upon arrival at the breeding site delayed clutch initiation, independent of arrival time. Extreme cold (sub-zero temperatures) within a staging area just prior to arrival at the breeding site carried over to prolong offspring development rate, potentially by influencing parental investment. Nest survival decreased with both later clutch initiation and prolonged offspring development, such that females that nested earlier and fledged offspring at a younger age were up to 45% more likely to reproduce successfully.
5. We demonstrate pronounced carry-over effects acting through mechanisms that influence breeding phenology and offspring development independently. We also highlight the potential importance of staging areas for alpine songbirds, particularly given that environmental conditions are becoming increasingly decoupled across seasons. Understanding the cross-seasonal mechanisms shaping breeding decisions in stochastic environments like the alpine enables more accurate predictions of future individual- and population-level responses to climate change.

## Introduction

Carry-over effects stem from conditions at one stage of the annual cycle that alter the state of an individual in subsequent stages, influencing behavioural decisions and fitness (Norris 2005, Norris and Marra 2007). Migratory songbirds spend up to 80% of the year in non-breeding habitats and may therefore be particularly susceptible to both time- and energy-based carry-over effects (Harrison et al. 2011, Marra et al. 2015). Delays in phenology can cascade across temporally-linked stages (Piersma 1987, Gow et al. 2019), while environmental factors that impact resource acquisition may reduce energy availability for reproduction or survival in subsequent stages (Gill et al. 2001, Paxton and Moore 2015, Rockwell et al. 2017). While related, time-based effects result in delayed phenology as individuals prioritize resource acquisition and survival over timing, while energy-based effects may occur when timing is critical to breeding success, forcing individuals to maintain phenology at sub-optimal energy levels (Descamps et al. 2011, Harrison et al. 2011). Linking breeding phenology and reproductive success to non-breeding habitat conditions allows us to identify critical periods in space or time throughout the annual cycle that influence individual fitness.

Non-breeding weather conditions or habitat quality can influence arrival times for individuals at the breeding site and subsequently clutch initiation date (Marra et al. 1998, Norris et al. 2003, Gunnarsson et al. 2005). In addition to a wintering site, songbirds can use stopover habitats for varying periods of time to accumulate the resources necessary for migration and breeding (Bayly et al. 2018, Briedis et al. 2018). Time-delays or energy-limitations may arise at each location, such that where and when songbirds use non-breeding habitats can influence the relative strength of carry-over effects (Marra et al. 2015). Nesting earlier is often linked to greater nesting success (Perrins 1970, Raquel et al. 2019), and fledging offspring with greater recruitment potential (Visser et al. 2015, Alves et al. 2019). Therefore, there should be strong selection for early clutch initiation (Lepage et al. 2000), and suboptimal conditions at any point in the annual cycle can theoretically produce negative fitness consequences through delayed breeding phenology.

For birds breeding in seasonal environments like the alpine, reproductive timing may be particularly critical to individual fitness (Bauer et al. 2016). Given short breeding seasons with limited opportunities to reproduce successfully, nesting early may be disproportionately beneficial (Martin and Wiebe 2004, Barras et al. 2021). However, stochastic habitats with frequent early-season storms may deplete energy reserves upon arrival, leading to decisions that prioritize survival over reproduction (i.e., nest abandonment; Martin et al. 2017, Wingfield et al. 2017). Under these conditions, selection on clutch initiation should be stabilizing (Coppack and Pulido 2009). While earlier clutch initiation can be risky, individuals arriving at the breeding site in better condition may invest more in parental care and benefit from early breeding even if conditions deteriorate (Bêty et al. 2003, Descamps et al. 2011). Energy-based carry-over effects may thus influence clutch initiation date in unpredictable habitats, but whether non-breeding conditions mitigate or compound the constraints of harsh, early-season conditions at the breeding site is unclear.

Offspring development reflects a potential link between clutch initiation dynamics and reproductive success. Nestling development—the period from hatch to fledge—is a critical, but vulnerable life-stage for altricial songbirds, where predation is the major source of fecundity loss and risk accumulates with time in the nest (Martin 1995, Martin and Briskie 2009). Nestlings that develop quickly can fledge earlier and potentially at a larger size, improving probability of survival in the first week outside of the nest when most post-fledging mortality occurs (Cox et al.2014, Martin et al. 2018). Rapid development may be constrained by early season storm events, prolonging time in the nest by restricting food availability and challenging thermoregulation (Stodola et al. 2010, Pérez et al. 2016, de Zwaan et al. 2020). The ability for parents to act as a buffer to these constraints and invest in maintaining nestling development rates likely depends on their individual condition (de Zwaan et al. 2019a). Carry-over effects may therefore influence offspring development and subsequently nest success through clutch initiation decisions or investment in parental care.

In an alpine population of horned lark (*Eremophila alpestris*) in northern British Columbia, Canada, age at fledging ranges from 7 to 13 days and is influenced by temperature, predation risk, and importantly, female body condition (de Zwaan et al. 2019a). Larks from this population are short-distance migrants (mean distance between breeding and winter site is 1080 km), yet some individuals remain at spring staging sites for prolonged periods that last up to 2 months (de Zwaan et al. 2019b). Therefore, spatially-structured environmental conditions during different time windows have the potential to inflict carry-over effects on breeding parameters.

Few studies assess the combined influence of weather conditions across non-breeding and breeding habitats on reproductive success, primarily focussing on breeding phenology (Mazerolle et al. 2011, Ockendon et al. 2013, Finch et al. 2014; but see Drake et al. 2014). Overall, the nature of carry-over effects stemming from different stages of the annual cycle and the critical time windows to which individuals respond are poorly understood, particularly for alpine birds.

Using 12 years of breeding data and 3 years of non-breeding location estimates from archival light-level geolocators (GLS), we assessed the influence of non-breeding and breeding weather conditions on clutch initiation date and nestling development rate. Specifically, we investigated: (i) the critical time windows over which temperature and precipitation variables may influence breeding phenology from three distinct regions (winter, staging, and local breeding sites), and (ii) whether non-breeding conditions influence nestling development rate, either through carry-over effects on clutch initiation (time-based) or independent of phenology (energy-based). Finally, (iii) we quantified the combined influence of clutch initiation date and nestling development rate on nest success.

## Methods

### Study system

Horned larks are open-country, ground-nesting songbirds (28–40 g) that breed in sparsely-vegetated habitat from 0 to over 4000 m above sea level (asl; Beason 2020). For 12 years (2003– 2007, 2010–2011, and 2015–2019), we studied a breeding population in ~3 km^2^ of alpine tundra (1650–2000 m asl) on Hudson Bay Mountain (HBM) in northern British Columbia (54.8°N, 127.3°W). Breeding conditions are challenging with high winds, fluctuating daily temperatures (–5 to +35 °C), and snow cover often extending into June (Camfield & Martin 2009, Martin et al. 2017). Breeding pairs initiate one to two nests per season with an average clutch of 3.6 eggs (range: 2–5; Camfield et al. 2010). Nestling development occurs primarily in June and early July (MacDonald et al. 2013). Females are responsible for most offspring care, building nests, incubating, and brooding without mate-feeding, while also provisioning nestlings more frequently than males (Goullaud et al. 2018).

At our site, larks are short-distance migrants that spend the winter east of the Cascade mountains in the south Columbia Plateau and the northern extent of the Great Basin region (Figure 1). During spring migration, they pass through the Okanagan Highlands, with some individuals making prolonged stopovers in this region from late-February to mid-April (average: 41 days, range = 21 – 66 days; de Zwaan et al. 2019b). Typical arrival date at the breeding site ranges between late-April and mid-May (de Zwaan et al. 2019b).

**Figure 1.**
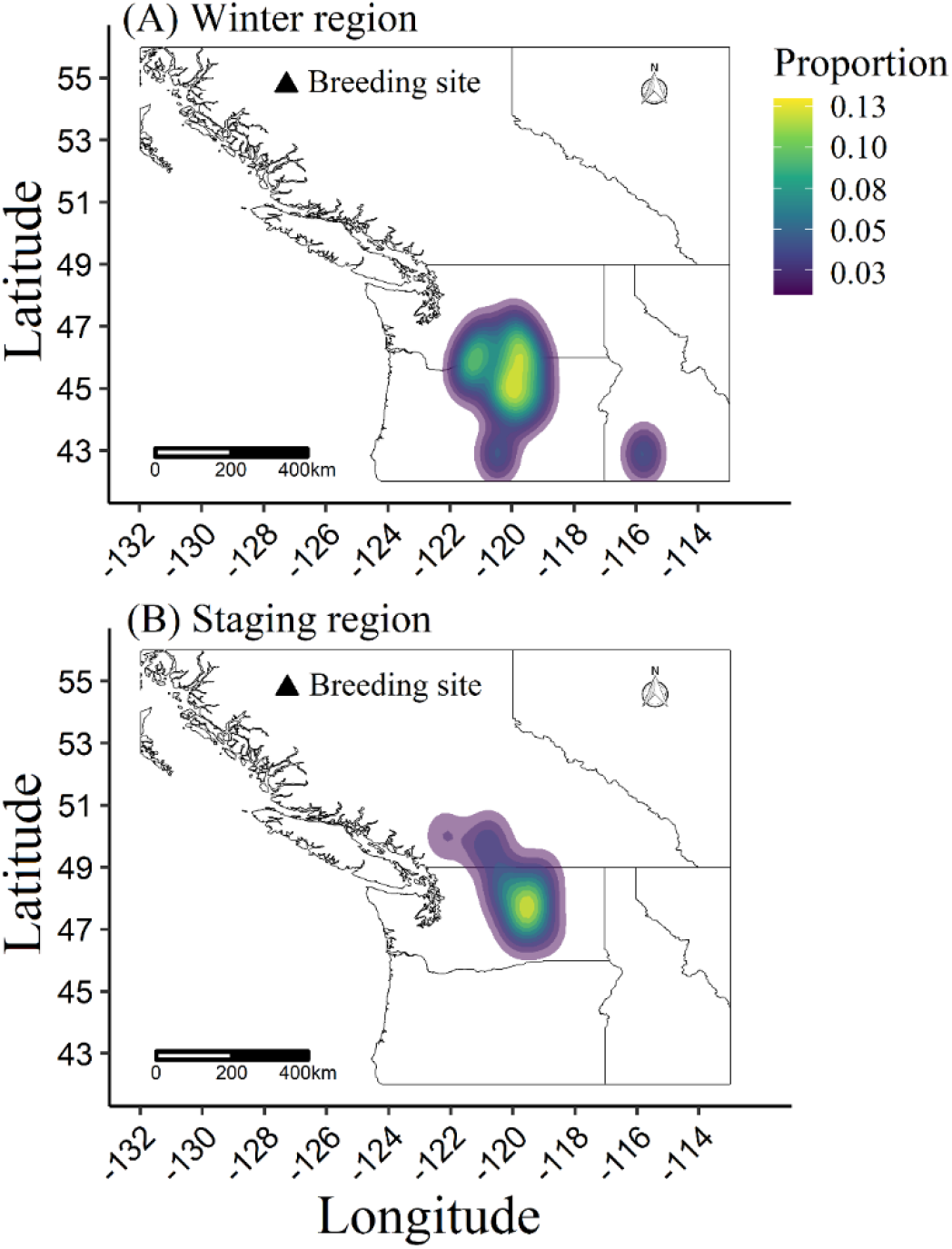
Kernel density plot showing the proportion of horned lark location estimates from de Zwaan et al (2019b) within the winter (A) and spring stopover (B) regions. Areas that contained at least 75% of the stage-specific location estimates were selected to differentiate the two regions such that the areas did not overlap. The black triangle denotes the breeding site.

### Field methods

Territories were monitored from before larks arrived at the breeding site each year until late-July. We located nests using a combination of systematic territory searches and behavioural cues. If a nest was located during the incubation stage, we back-calculated clutch initiation date from hatch date using an average incubation period of 12 days and assuming that incubation was initiated on the penultimate egg (Camfield et al. 2010). Nests were monitored every 2–3 days until 10 days of incubation or 5 days post-hatch after which we visited nests daily to determine hatch and fledge date. Age at fledging was calculated as the number of days between hatch and fledge (hatch date = 0). Adults were captured at the nest using either a bownet or noose-line trap and banded with a unique combination of one USGS aluminum band and three colour bands for subsequent identification (de Zwaan et al. 2018). All nests were classified as first nests, re-nesting attempts following failure, or double-brooding attempts.

### Local weather variables

Hourly precipitation and temperature variables at the breeding site were recorded using three HOBO weather stations (Onset Computer Co., Pocaset, MA, USA): 1) a Micro Station Logger H21-002 from 2005 to 2007, 2) a U30-NRC station for 2010 to 2017, and 3) an RX3000 satellite station for 2018 and 2019. While the latter two stations were positioned ~1.2 km west of the original station, all three were located at 1,695 m asl. The sensors were placed 1.5 and 3 m above ground, and less than 2.4 or 1.2 km from any nest for 2005–2007 and 2010–2019, respectively.

Precipitation values were missing for 2003, 2004, and part of 2017. To estimate precipitation, we used values from the Smithers Regional Airport (522 m asl; ~8 km from study site). Daily precipitation varied between the airport and study site, but consistently identified days with moderate precipitation (≥ 1 mm; 81% accuracy; Martin et al. 2017). Therefore, to remain consistent, we classified each day into a binomial ‘precipitation day’ (precipitation = 1; no precipitation = 0) whether we were using data from the Smithers Airport or the weather station at the study site. For temperature values which were also missing in 2017, we interpolated hourly estimates of air surface temperature from the 8 nearest grid points in the National Centers for Environmental Prediction (NCEP) R-1 dataset using the R package “RNCEP” (Kemp et al. 2012). To validate this method, we compared the interpolated estimates to measured temperature from our study site for the 11 years with existing data. Estimates and true measurements were highly correlated (r_p_ = 0.91), but the intercept differed, likely due to an elevation effect. We therefore subtracted the value of the intercept difference (1.78°C) from all interpolated estimates to better approximate true temperature measurements.

We retrieved snow depth data for 2009–2019 from the HBM ski resort monitoring station. From 2003–2007, we extracted data from the nearest weather stations involved in the automated snow weather station array of the Provincial Snow Survey Network (B.C. Ministry of Environment and Climate Change Strategy 2019). Snow depth data from Shedin Creek station (120.5 km north; 1,480 m asl) correlated strongly with HBM data from 2009–2019 (r_p_ = 0.90), and therefore we used these data as a proxy of snow depth at our study site from 2003–2007. We evaluated snow depth on April 15^th^ because this is the latest date with consistent data across years and, being just prior to when larks arrive in the alpine, is likely to influence breeding decisions (de Zwaan et al. 2019b).

### Non-breeding weather variables

From 2016 to 2018 we retrieved 17 light-level geolocators (Intigeo-P65B1-11; Migrate Technology Inc.), representing 9 females and 8 males. For full details on calibration and location estimation, see de Zwaan et al (2019b). Briefly, we used the “Hill-Ekstrom” calibration method to calculate a zenith angle from stationary periods near the equinox (Hill & Braun 2001, Ekstrom 2004). Non-breeding location estimates were then derived by locating likely stopovers of ≥ 3 days within a distance threshold of 150 km, and then fitting a movement model with a prior probability distribution for flight speed and a land mask that restricted individuals to stopover in open-country habitat only. Location estimate error averaged ± 58 and 61 km for latitude and longitude, respectively (de Zwaan et al. 2019b).

From these location estimates, we created kernel density plots to quantify winter and staging regions (Figure 1). Most of the location estimates could be classified into two distinct areas: 1) winter region (Latitudinal range: 44–47°N; Longitudinal range: 121.5–119°W), and 2) staging area (47–49°N; 120.5–118.5°W). While there was some peripheral overlap, explicitly separating the stopover and winter locations allowed us to test if weather conditions closer to the breeding site, both in time and space, may influence breeding parameters.

We gathered temperature and precipitation data separately for the winter and staging areas. Within each region, we laid out a grid of points spaced at 50 km intervals. We then interpolated surface air temperature (‘air.sig995’) and precipitation rate (‘prate.sfc’) from the NCEP R-1 database to each point four times daily (midnight, 0600, 1200, and 1800 hr) for all 12 years of the study. The daily average, min, and max temperatures were calculated per point and then averaged over the region for each day. If the regional average daily temperature was ≤ 0° C, we denoted this as a ‘freeze day’. Similarly, we converted the point estimate precipitation rate to total mm/m^2^ per day and averaged within each region for mean daily precipitation.

Overall, for the non-breeding regions (winter, staging) and the breeding site, temperature variables consisted of daily average, minimum, and maximum temperatures, as well as freeze days (≤ 0° C). For precipitation, we considered average daily precipitation (mm/m^2^) for the non-breeding regions and precipitation days (≥1mm) for the breeding site.

### Statistical analysis

#### Weather variable selection and model fitting

We assessed the influence of spatially-explicit weather conditions on clutch initiation and age at fledging for first nests only. For each response variable, we used a sliding window approach within the R package ‘climwin’ (Bailey and van de Pol 2015) to identify the most influential weather variables and associated time windows for the winter region, staging area, and breeding site. This systematic approach tests the influence of each weather variable across all possible time windows and then ranks each model with Akaike Information Criterion for small sample sizes (AICc; van de Pol et al. 2016). We then extracted the top time windows for each weather variable to build a global candidate model and used model averaging to estimate effect size and variable importance. All analyses were performed using R version 3.6.3 (R Core Team 2020).

We constructed a sliding window that spanned different but overlapping time periods for each of the three regions of interest based on prior knowledge of when individual larks arrive and depart from each region. These time windows consisted of December 1 to May 1, February 15 to May 1, and April 15 to June 1 (clutch initiation) or June 12 (age at fledging) for the winter, staging, and breeding sites, respectively. These time periods captured when individuals were most likely to be at the wintering, staging, and breeding sites (de Zwaan et al. 2019b), while June 1 and June 12 were chosen as reference points as they represent the average clutch initiation and hatch date for first nests across all years. We constrained the minimum time window to 7 days for winter and staging conditions, and 3 days for the breeding site, with the maximum window size allowable being the entire time window. The minimum non-breeding time windows were chosen to better reflect weather patterns rather than anomalies and to avoid spurious correlations with short time periods that do not make ecological sense. A minimum window of 3 days was selected for the breeding site to allow time for female larks to exhibit a physiological response (i.e., ova development) and make reproductive decisions based on prevailing weather conditions (Williams 2012, de Zwaan et al. 2020).

For each weather variable, all possible time windows were ranked with AICc relative to the null model (year as the only predictor). The ‘best’ time window was chosen based on the lowest AICc if it had a significantly better fit than the null (ΔAICc < –2). If multiple time windows were within 2 AICc of the top model, we chose the one with the largest β-coefficient. Multiple comparisons can lead to spurious results, so we used the built-in randomization procedure in ‘climwin’ to determine the likelihood of selecting the same top model by chance (Type 1 error). Each model was run on 100 randomized datasets and the ΔAICc of the observed data was compared to the ΔAICc of each randomization. Weather variables where the observed results were significantly different (P < 0.05) from the randomized results were selected as candidate variables.

Since larks from this population are short-distance migrants, temperature and precipitation values for different regions and time periods may be spatially or temporally autocorrelated (Legendre and Fortin 1989). We used Pearson’s product-moment correlations and Variance Inflation Factors (VIF) to test for possible multicollinearity among candidate weather variables. For clutch initiation, winter temperature and precipitation were strongly correlated (r_p_ = –0.80; Supplemental Appendix: Figure S1). Therefore, we excluded models containing both variables from consideration when conducting AICc model selection and averaging (see below). For age at fledging, winter and stopover freeze days were strongly correlated (r_p_ = 0.87; Supplemental Appendix: Figure S2). These variables represented similar time windows (winter = Mar 27 – Apr 17; stopover = Apr 1 – Apr 22) and were likely capturing the same weather pattern. Therefore, we chose to only include stopover freeze days because it had the greatest effect size and lowest AICc relative to the null during sliding window selection (stopover ΔAICc = –7.4; winter ΔAICc = –4.5). Following these steps, all predictor variables of both clutch initiation and age at fledging had a subsequent VIF < 1.9, indicating no multicollinearity.

We built global models for both clutch initiation and age at fledging which included the selected candidate weather variables and additional biologically relevant explanatory variables. For clutch initiation date, we included snow depth as an explanatory variable. For age at fledging, we included clutch initiation date and brood size to control for potential effects on development rate. For both response variables, we also included year as a linear term to test for an overall trend over the study period. All variables were standardized and centered to allow comparison of relative effect sizes. We broke each global model into all possible subset models (clutch initiation: n = 64 models; age at fledging: n = 32) and ranked each with AICc. As mentioned above, clutch initiation sub-models that included both winter temperature and precipitation were not considered, allowing us to determine which winter weather variable best fit the data. Each sub-model was bootstrapped 1000 times and models with Akaike weights summing to 0.90 were averaged to produce the final coefficients (Burnham and Anderson 2002; see Supplemental Appendix: Table S1 and S2 for top models). Bootstrapped means and 85% credible intervals were used to evaluate parameter strength and significance. We chose an 85% CI because it is more consistent with an information theoretic approach (Arnold 2010).

### Fitness effects

We assessed the influence of advanced or delayed clutch initiation relative to the annual mean on daily nest survival (DNS). We centered clutch initiation within year such that negative values were earlier than average and positive values were later. We fit a mixed-effects discrete logistic exposure model with a complementary log-log link to model the probability of successfully fledging offspring while accounting for exposure days. Year was included as the random effect and relative clutch initiation as the only fixed effect. We fit two separate models, with clutch initiation as a linear and quadratic term. A linear relationship would be expected if nest survival was highest early in the season and declined with greater predator activity as the season progressed. Conversely, nest survival could increase with time as less snow cover and a greater density of bird nests may produce a dilution effect with more potential nest sites, decreasing the likelihood of predators locating an individual nest (Chalfoun and Martin 2009). A quadratic relationship would occur if nest survival was highest for early and late nests because predators were targeting nests during peak nesting activity at some middle date (Wilson et al. 2007). Model support was compared using AICc.

## Results

Over 12 years, 382 first nests were monitored. Average clutch initiation date was June 1, spanning only 11 days across years from May 28 to June 8 (Figure 2A). Variation among first nests of the same year ranged from 10 to 26 days between the first and last clutch initiated (mean = 17.3 days). For age at fledging, we could only use 10 years of data, as 2007 had no successful first nests and 2010 had one (see Supplemental Appendix: Table S4 for annual sample sizes). For 118 nests, age at fledging ranged from 7 to 13 days. Average age at fledging across all years was 9.2 ± 0.1 (SE) days, with a minimum within-year average of 8.4 and maximum of 10.2 days (Figure 2B).

**Figure 2.**
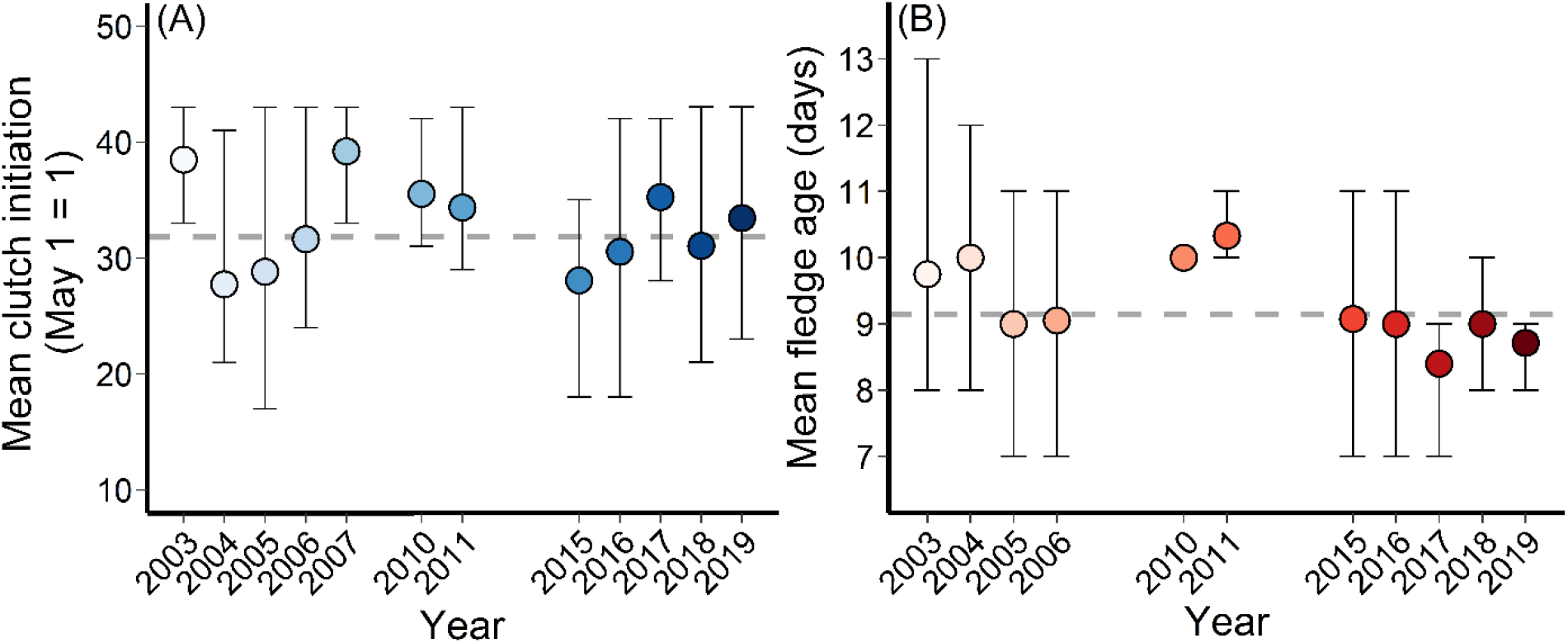
Within-year averages of (A) clutch initiation date and (B) age at fledging for an alpine breeding population of horned lark in northern British Columbia, Canada. Error bars depict the range of values (min, max) and the dashed grey line is the among-year average. In (B), 2010 represents only one nest and thus was not included in analysis but is depicted here for visualization purposes. The colour gradients are used to visualize progression over time.

### Clutch initiation date

Sliding window analysis identified winter temperature (best window: Dec 2 – Mar 10), winter precipitation (Feb 10 – Apr 10), and stopover precipitation (Feb 21 – Apr 9) as the most likely non-breeding predictors of clutch initiation date. Local breeding temperature (May 1 – May 22) and precipitation (May 1 – May 9) were selected as the most likely breeding site correlates.

After model selection, winter precipitation was a better predictor of clutch initiation than winter temperature, which did not factor into any of the top models (Supplemental Appendix: Table S1). Model averaged estimates indicated that greater winter precipitation delayed clutch initiation (β = 0.25; Figure 3A). For every 0.5 mm/m^2^ increase in average daily winter precipitation, clutch initiation date was delayed by 4.5 days (Figure 4A). At the breeding site, higher temperatures prior to breeding were linked to earlier egg laying dates (β = –0.41; Figure 3A). For every 1°C increase in average temperature during the month of May at the breeding site, clutch initiation date advanced by 1.5 days (Figure 4B). Stopover and local precipitation, as well as snow depth failed to influence clutch initiation date, and there was no overall annual trend (Figure 3A; Supplemental Appendix: Table S3).

**Figure 3.**
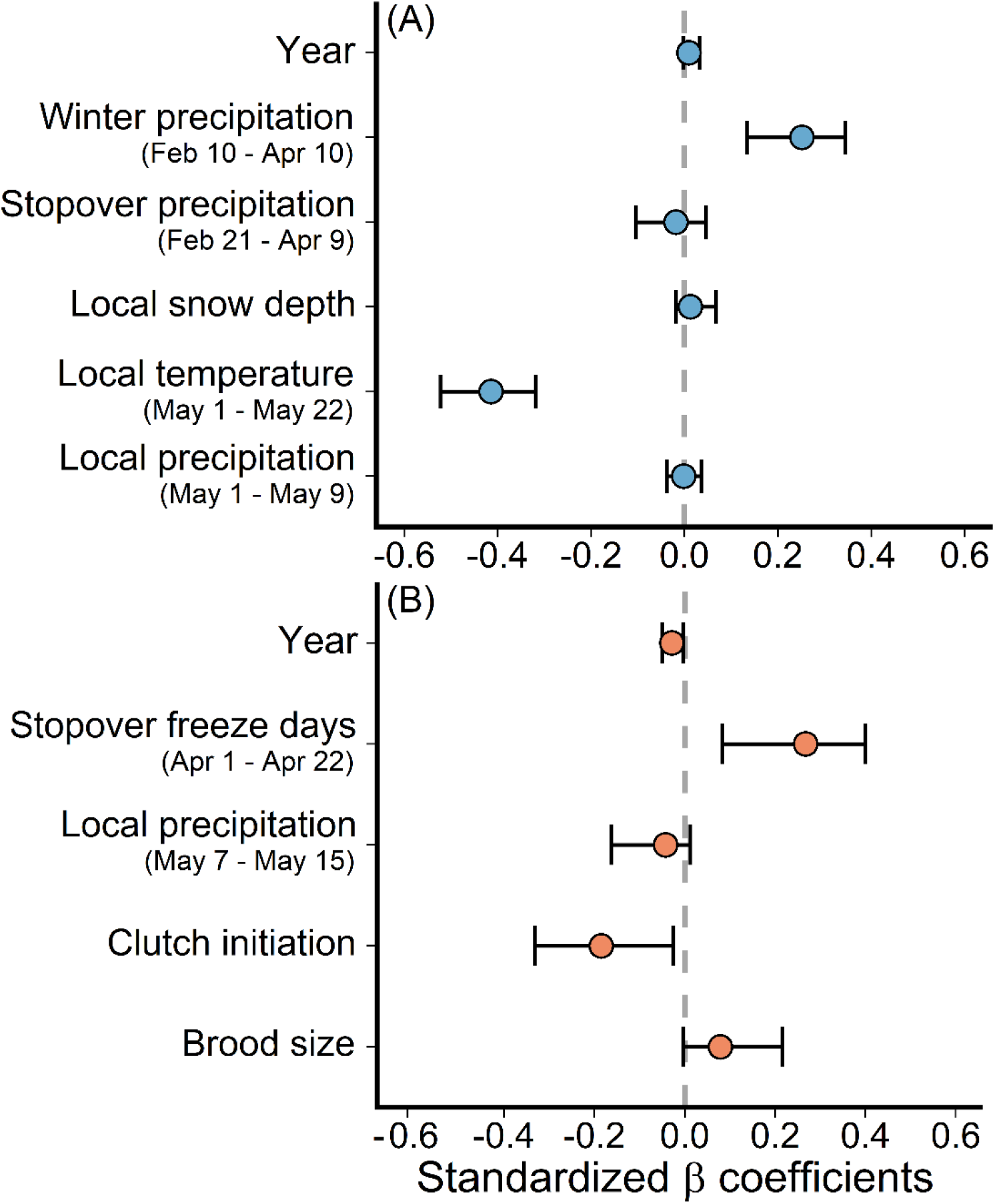
Standardized β-coefficients and 85% credible intervals as estimated from model averaging and 1000 bootstraps for (A) clutch initiation and (B) age at fledging. The points depict the bootstrapped means and the error bars are the 85% credible intervals. Variables are considered significant if their 85% CI does not include zero (the grey dashed line) to be more consistent with AICc model selection. ‘Local’ refers to the breeding site.

**Figure 4.**
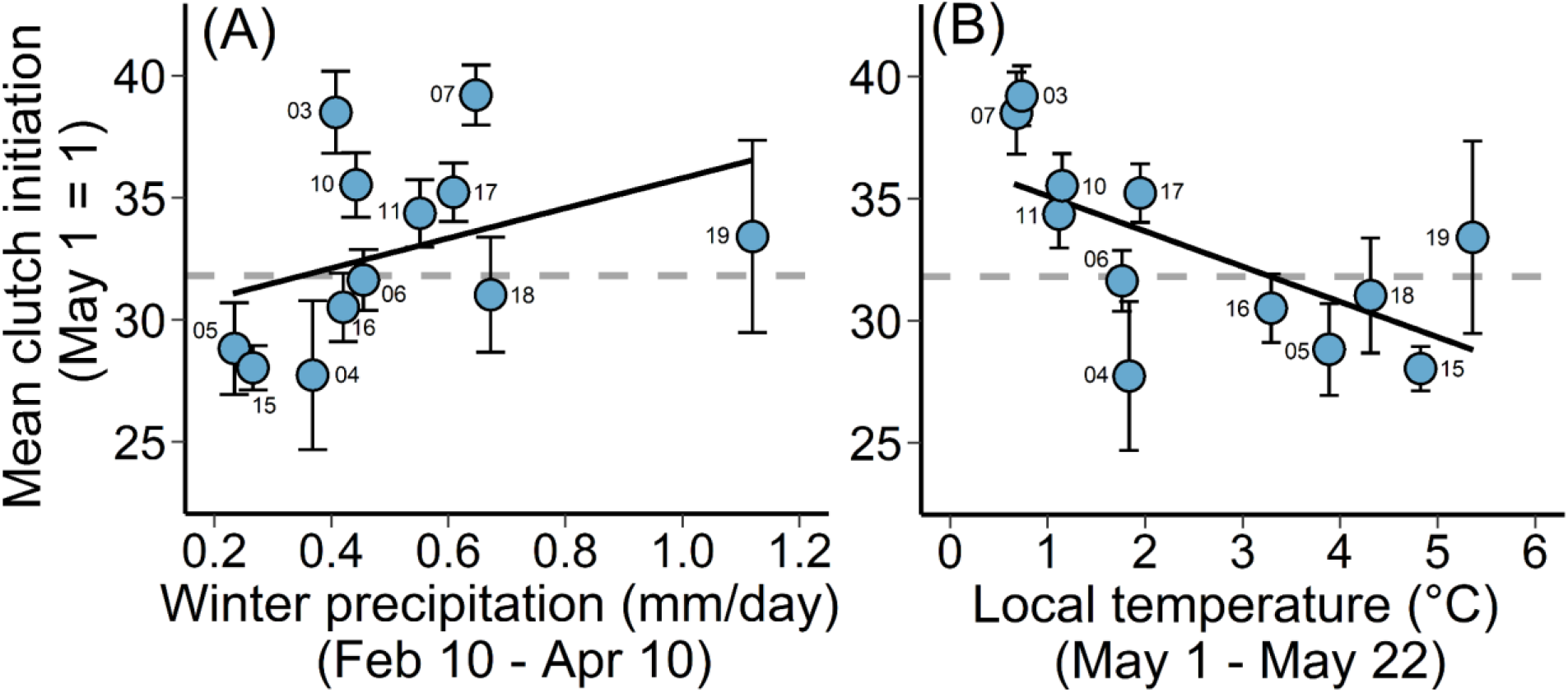
The influence of (A) winter precipitation and (B) local temperature on average clutch initiation date among years. The dashed grey line represents the average across all years, while the points and error bars are the within-year averages and standard errors, respectively, labelled by year (last two digits). The trend lines are derived from the bootstrapped models.

### Age at fledging

The number of days below freezing at the staging site (i.e., freeze days; Apr 1 – Apr 22) and precipitation at the breeding site (local precipitation; May 7 – May 15) were the only weather variables selected with the sliding window approach for age at fledging. After model averaging, a greater proportion of freeze days at the staging area was associated with prolonged nestling development (i.e., older age at fledging; β = 0.26; Figure 3). For every 2 days below 0°C in April, age at fledging increased by 0.15 days (Figure 5). Females that initiated nests later fledged offspring significantly sooner (β = –0.18; Figure 3B), and there was a slight negative trend in average age at fledge between 2003 and 2019 (β = –0.03; Figure 3B). Local precipitation and brood size did not influence age at fledge (Figure 3B; Supplemental Appendix: Table S3).

**Figure 5.**
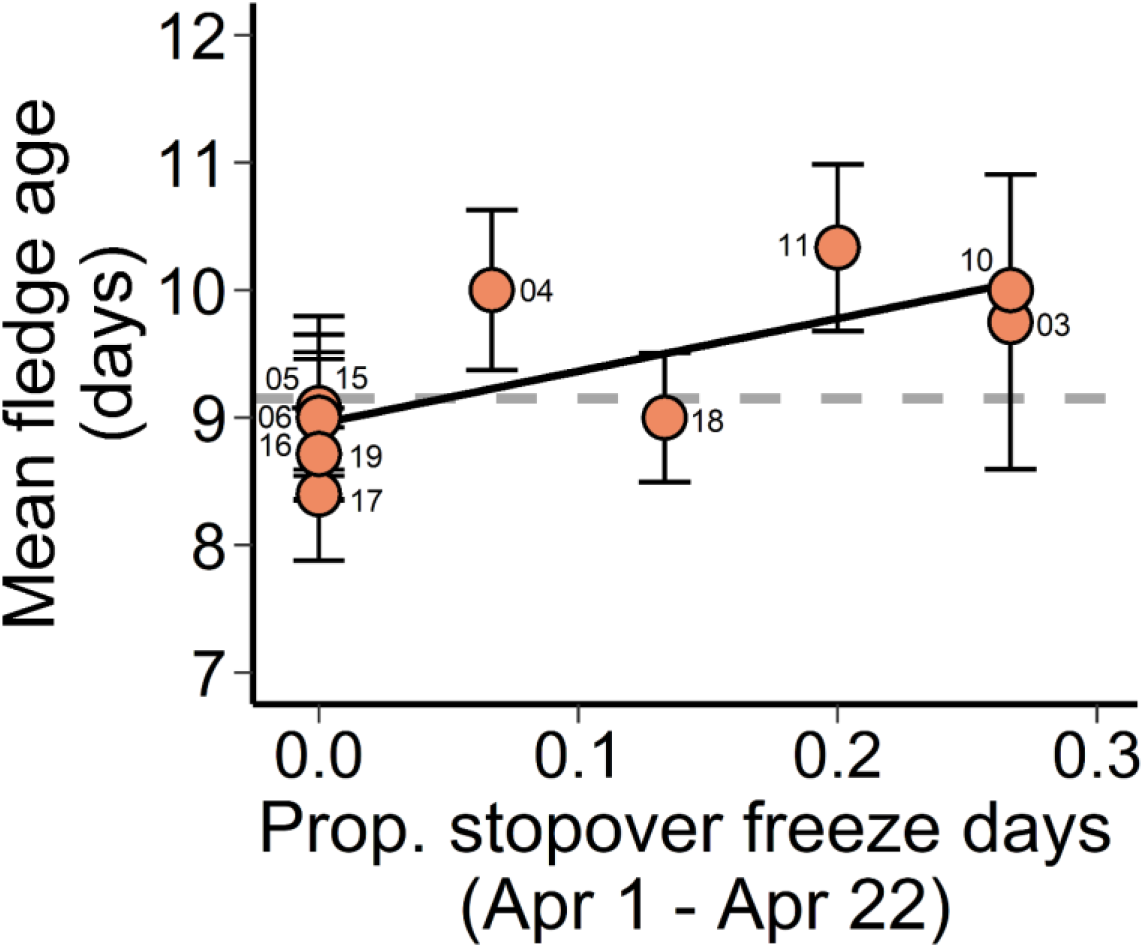
The relationship between the proportion of stopover freeze days (days ≤ 0°C) and average age at fledging. The dashed grey line represents the average across all years, while the points and error bars are the within-year averages and standard errors, respectively, labelled by year (last two digits). The trend lines are derived from the bootstrapped models. For 2010, a point is included for reference, but the datum was not involved in the analysis due to a low sample size (n = 1) and thus did not influence the trend line.

### Fitness consequences

The linear and quadratic daily nest survival (DNS) models did not differ (ΔAICc ≤ 2) and therefore the linear relationship was selected as the most parsimonious. For a mean clutch initiation date, the average estimated DNS across years was 0.967 ± 0.002 (± SE), corresponding to a 44.2 ± 1.2% probability of successfully fledging offspring over a 24-day nest cycle. This ranged from a DNS low of 0.492 ± 0.281 in 2007 to a high of 0.997 ± 0.002 in 2004 (Supplemental Appendix: Table S4).

Within years, DNS declined with clutch initiation date (z = –2.85, P < 0.01). For the earliest observed clutch initiation (13 days earlier than average), the predicted DNS was 0.986 ± 0.003 or 71.0 ± 3.8% nest success, compared to 0.940 ± 0.017 or a 22.8 ± 4.8% nest success for the latest observed clutch initiation (14 days later than average). The fitness consequences of nestling development time were dependent on clutch initiation date. For an average clutch initiation, prolonged development times (13 days) had a predicted 14.6% lower probability of success than rapid development (7 days; Figure 6). For early nests, the predicted cost of delayed fledge was minimal (–6.9%), but the cost increased as the season progressed (–20.1%; Figure 6).

**Figure 6.**
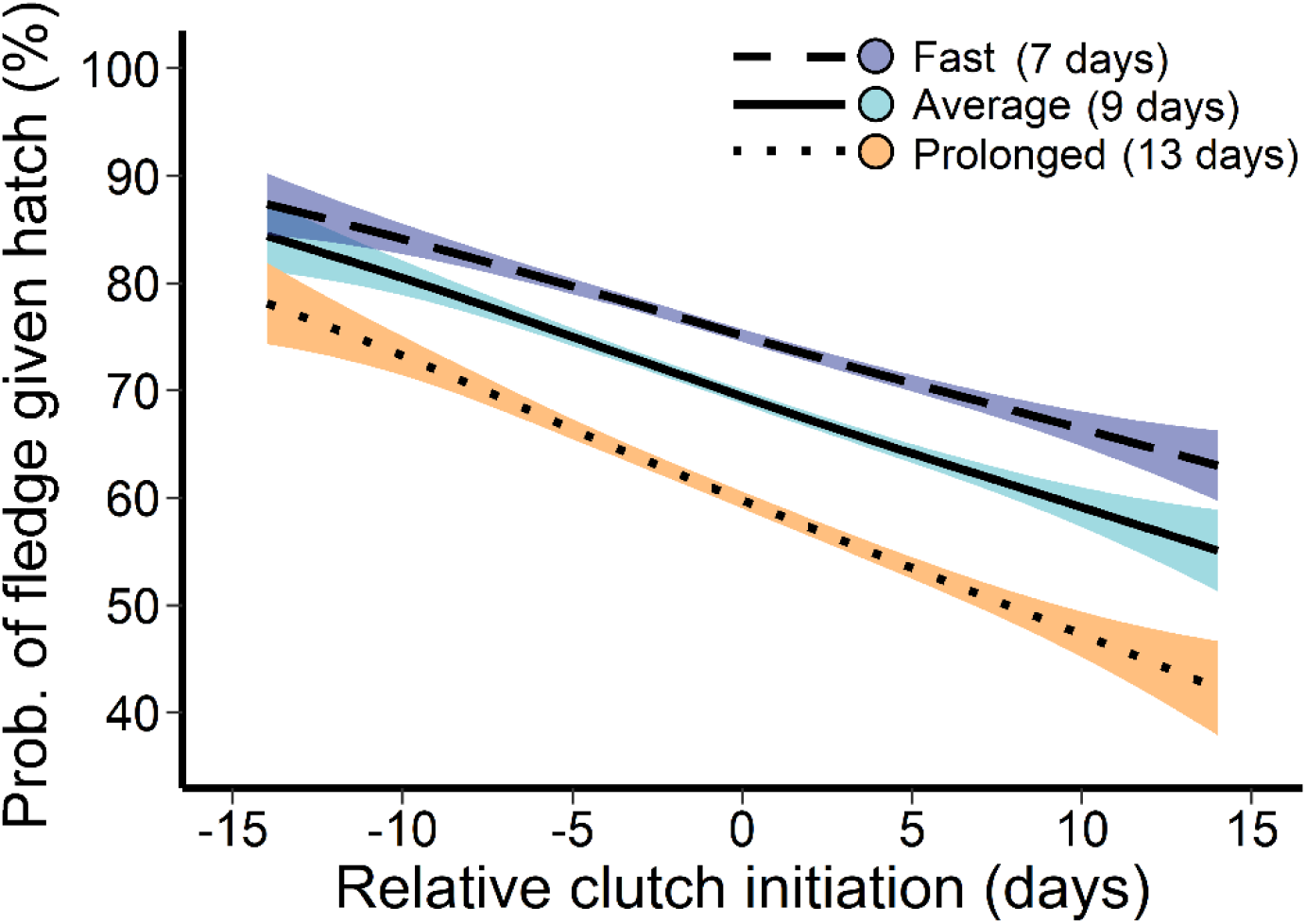
The association between clutch initiation date and probability of fledging given hatch (i.e., nestling survival) across minimum (7 days), average (9.2 days), and maximum (13 days) observed nestling development times. Slopes are predicted from the relationship between clutch initiation and average daily nest survival (DNS) based on the observed within-year range of clutch initiation dates (0 = within-year average). The shaded bands depict the 95% CI of the bootstrapped residuals.

## Discussion

We demonstrate that among year variation in clutch initiation date and nestling development time is influenced by strong carry-over effects stemming from spatially explicit weather conditions across the annual cycle. Periods of extreme cold at spring stopover sites prolonged nestling development time independent of an effect on clutch initiation date. Given this independence and that arrival date at the breeding site does not influence clutch initiation in this population (de Zwaan et al. 2019b), our results align most with expectations for energy-rather than time-based carry-over effects (Harrison et al. 2011). In addition, while later clutch initiation is often linked to reduced nest success and fecundity (Rockwell et al. 2012, Imlay et al. 2018), we show that delayed clutch initiation and prolonged offspring development combine synergistically to reduce nest success. Therefore, suboptimal conditions at critical points in the non-breeding season can influence individual fitness through carry-over effects on both clutch initiation and nestling development.

### Clutch initiation

Colder, early season temperatures at the breeding site, but not snow depth, delayed clutch initiation. Larks are early breeders and can build nests as soon as small patches of ground are snow-free (Beason 2020). Snow depth may therefore only influence clutch initiation at a more localized scale because topography and temperature can combine to drive heterogeneity in snowmelt, exposing nesting habitat independent of average snow depth (e.g., on exposed ridges; Niittynen et al. 2018). Even without snow as a constraint, synchronizing hatch date with resource availability can be critical to nest success (Visser et al. 2006). Warmer spring temperatures at the breeding site are often associated with earlier clutch initiation (Bowers et al. 2016, Imlay et al. 2018, Ram et al. 2018). In the alpine, foliage leaf-out and insect abundance increase with snowmelt and warmer temperatures over the breeding season in an extended resource ‘wave’ (García-González et al. 2016). Therefore, postponing clutch initiation with colder early-season temperatures may represent resource tracking by female larks, although this decision likely depends on individual condition and predation risk costs as the season progresses (de Zwaan et al. 2019a).

While greater than average precipitation at the winter site delayed clutch initiation, colder temperature and greater precipitation were highly correlated, potentially indicating the influence of harsher winter conditions on breeding phenology. Short-distance migrants that breed in naturally variable habitats such as the alpine may be more flexible in their breeding phenology than long-distance migrants (Boelman et al. 2017, Mizel et al. 2017), and therefore time-based carry-over effects may be less pronounced (Briedis et al. 2018, Gow et al. 2019). Instead, harsh winter conditions may disrupt resource acquisition and influence individual body condition upon arrival at the breeding site, requiring females to recover energy stores prior to breeding (Bêty et al. 2003, 2004). In alpine habitats, challenging early season conditions may compound the effects of harsher winters, prolonging the time needed to invest in self-maintenance (Harrison et al. 2013). Intuitively, this result also highlights the potential for more favourable breeding site conditions to moderate the influence of poor winter conditions (Descamps et al. 2011).

### Age at fledge

Periods of sub-zero temperatures in April at the staging grounds increased average age at fledging. A similar relationship was not observed for average temperature, suggesting that only severe conditions induced cross-seasonal effects on nestling development. While most individuals pass through the stopover region during northward migration, the time spent in this area can range from days to months (de Zwaan et al. 2019b) and therefore exposure to staging conditions varies among individuals. However, 10 of 17 tracked larks (59%) remained within the staging area ≥ 21 days and experienced improved reproductive success, producing an average of 1.8 more fledglings than individuals that did not stop in the staging area (de Zwaan et al. 2019b). Given the prolonged duration of stay and the association with both reproductive success and nestling development rate (this study), the Okanagan Highlands staging region could be an important component of the annual cycle, particularly in April.

Staging conditions did not influence clutch initiation, indicating that prolonged development time is not a result of cascading time-dependencies. Instead, periods of extreme cold may induce changes in female physiology or body condition shortly before breeding (Sorenson et al. 2016). Additionally, poor body condition prior to breeding can reduce investment in parental care behaviours such as incubation and offspring provisioning in favour of self-maintenance (Williams 2012). Resource-challenged females may limit incubation bouts, leading to cooled embryos (MacDonald et al. 2013, Coe et al. 2015) and reduced nestling development rates (Nord and Nilsson 2011). Interestingly, colder temperatures during the egg laying stage have been linked to faster nestling development in horned lark (de Zwaan et al. 2020), highlighting further that both the timing and severity of weather events can have important, but sometimes contrasting implications for breeding performance. While time-series of female physiology and body condition are necessary to determine the underlying mechanisms, we demonstrate that conditions during migration can influence offspring development independent of effects on clutch initiation date.

### Fitness effects

Carry-over effects originating across the annual cycle that influence clutch initiation and nestling development can combine synergistically to impact individual fitness. Nest survival of first nests declined linearly with time of season. Prolonged development time had minimal negative consequences for early nests, but resulted in drastically reduced success of later nests. With declining nest success, the association between later clutch initiation date and faster offspring development rate potentially indicates a compensatory response to delayed start that may be facilitated by increasing food availability. Notably, slower development is hypothesized to be adaptive under stochastic conditions (Arendt 1997), with rapid development potentially leading to physiological constraints on fitness later in life (Monaghan 2008, Criscuolo et al. 2011).

Therefore, females able to initiate clutches earlier may benefit from disproportionately greater fitness through a combination of reduced nest predation risk and longer offspring development periods resulting in higher quality offspring. Early-breeding female larks in good body condition fledge larger offspring at an earlier age than poor condition females (de Zwaan et al. 2019a), indicating that it is possible to balance the benefits and constraints of nesting early. While early clutch initiation is frequently linked to greater reproductive success, the underlying mechanisms are often unclear (Morrison et al. 2019). We propose that offspring development is a component of this association.

## Conclusion

Early clutch initiation and rapid offspring development can be adaptive, but environmental conditions throughout the annual cycle may constrain the ability to advance phenology. For an alpine population of larks, we demonstrate that both non-breeding carry-over effects and breeding site conditions can influence reproductive phenology as well as nestling development rate with direct fitness consequences. We highlight that carry-over effects may be particularly pronounced for birds breeding in fluctuating habitats like the alpine, where stochastic early season conditions and resource availability coupled with a short breeding season limit reproductive opportunities. The climate is changing faster at higher latitudes and elevations (Pepin and Lundquist 2008, Zhang et al. 2019), potentially decoupling conditions between non-breeding and breeding habitats for alpine birds and limiting the capacity for flexibility in breeding phenology through unreliable environmental cues (Charmantier and Gienapp 2014, Senner et al. 2018). By addressing the influence of conditions across the annual cycle, we can better understand the mechanisms driving breeding decisions in stochastic habitats and make predictions about future individual- and population-level responses to climate change.

## Acknowledgements

We thank C. Ames, A. Clason, M. Martin, M. Mossop, L. Sampson, M. Tomlinson, L. Wenn, M. Grabowski, C. Storey, A. Sulemanji, D. Maucieri, N. Froese, C. Rivas, N. Bennett, E. Gow, S. Hudson, T. Altamirano, and N. Morrel for their contributions to data collection. Special thanks to A. Chalfoun, C. Beckmann, and S. Wilson for providing feedback on preliminary drafts. Field work was conducted on the traditional unceded territory of the Wet’suwet’en nation.

## Funding

Funding for this research was provided to DRD by the Northern Scientific Training Program (NSTP), American Ornithological Society (AOS), Society of Canadian Ornithologists, and Hesse Research Award, to AFC, ECM, and KM by the Natural Sciences and Engineering Research Council of Canada (NSERC), NSTP, and Environment and Climate Change Canada (ECCC), and to AFC by the AOS. Financial support was provided to DRD by NSERC and the University of British Columbia.

## Ethical statement

Data were collected under permit from Environment and Climate Change Canada and the University of British Columbia IACUC protocols A03-0095, A07-0048, and A15-0027.

## Authors’ Contributions

DRD and AD conceived the ideas; DRD, AFC, ECM, and KM designed the methodology and collected the data; DRD analysed the data and led writing of the manuscript. All authors contributed critically to the drafts and gave final publication approval.

## Data Accessibility

Data and code will be made available in the Figshare public repository.

## Conflict of Interest

The authors declare no conflict of interest.

## Supplementary Appendix

**Table A1.** Top candidate models for clutch initiation date ranked by AICc prior to bootstrapping and model averaging. Local temp and precip refers to temperature and precipitation for the breeding site. Of the total 64 models, only those with Akaike weights that sum to 0.90 are displayed. Values below the model terms are the standardized β coefficients.

| Local temp | Local precip | Snow depth | Staging precip | Winter precip | Year | df | AICc | ΔAICc | Akaike weight |
| --- | --- | --- | --- | --- | --- | --- | --- | --- | --- |
| -0.38 | — | — | — | 0.26 | — | 4 | 985.2 | 0.00 | 0.24 |
| -0.42 | — | — | — | 0.23 | 0.01 | 5 | 986.1 | 0.93 | 0.16 |
| -0.38 | — | 0.03 | — | 0.27 | — | 5 | 986.8 | 1.64 | 0.11 |
| -0.39 | — | — | -0.04 | 0.28 | — | 5 | 986.9 | 1.67 | 0.11 |
| -0.38 | 0.01 | — | — | 0.26 | — | 5 | 987.3 | 2.05 | 0.09 |
| -0.42 | — | 0.03 | — | 0.24 | 0.01 | 6 | 987.8 | 2.58 | 0.07 |
| -0.43 | — | — | -0.03 | 0.25 | 0.01 | 6 | 987.9 | 2.65 | 0.07 |
| -0.43 | -0.02 | — | — | 0.23 | 0.01 | 6 | 988.1 | 2.91 | 0.06 |
| -0.38 | — | 0.02 | -0.02 | 0.28 | — | 6 | 988.8 | 3.58 | 0.04 |
| -0.37 | 0.02 | 0.03 | — | 0.27 | — | 6 | 988.8 | 3.62 | 0.04 |

**Table A2.** Top candidate models for age at fledging ranked by AICc prior to bootstrapping and model averaging. Local precip refers to precipitation recorded at the breeding site. Of the total 16 models, only those with Akaike weights that sum to 0.90 are displayed. Values below the model terms are the standardized β coefficients.

| Stopover<br>freeze days | Local<br>precip | Clutch<br>initiation date | Brood<br>size | Year | df | AICc | ΔAICc | Akaike<br>weight |
| --- | --- | --- | --- | --- | --- | --- | --- | --- |
| 0.28 | – | -0.20 | – | -0.03 | 5 | 326.5 | 0.00 | 0.25 |
| 0.28 | – | -0.22 | 0.11 | -0.03 | 6 | 327.1 | 0.61 | 0.18 |
| 0.26 | -0.07 | -0.18 | – | -0.03 | 6 | 328.2 | 1.66 | 0.11 |
| 0.26 | -0.06 | -0.19 | 0.10 | -0.03 | 7 | 329.0 | 2.46 | 0.07 |
| 0.31 | – | -0.18 | – | – | 4 | 329.1 | 2.55 | 0.07 |
| 0.21 | -0.14 | – | – | -0.03 | 5 | 329.3 | 2.78 | 0.06 |
| 0.31 | – | -0.20 | 0.12 | – | 5 | 329.4 | 2.90 | 0.06 |
| 0.24 | – | – | – | -0.03 | 4 | 329.5 | 3.02 | 0.05 |
| 0.29 | -0.09 | -0.15 | – | – | 5 | 330.4 | 3.88 | 0.04 |
| 0.24 | -0.15 | – | – | – | 4 | 330.6 | 4.07 | 0.03 |
| 0.21 | -0.14 | – | 0.08 | -0.03 | 6 | 330.8 | 4.23 | 0.03 |
| 0.24 | – | – | 0.08 | -0.03 | 5 | 330.9 | 4.37 | 0.03 |
| 0.29 | -0.08 | -0.17 | 0.11 | – | 6 | 331.0 | 4.48 | 0.03 |

**Table A3.** Model outputs for clutch initiation and age at fledging, depicting the mean bootstrapped standardized coefficients, standard error (SE), and 85% credible intervals (CI). Variable importance (VI) is the sum of the Akaike weights for all models that include that parameter. Significant predictors (85% CI does not include zero) are in bold.

| | Mean<br>β-coefficient | SE | Lower CI | Upper CI | VI |
| --- | --- | --- | --- | --- | --- |
| *Clutch initiation* |  |  |  |  |  |
| Year | 0.01 | 0.01 | −0.01 | 0.03 | 0.39 |
| **Winter precipitation** | 0.25 | 0.07 | 0.14 | 0.34 | 1.00 |
| Stopover precipitation | −0.02 | 0.07 | −0.11 | 0.04 | 0.30 |
| Snow depth | 0.02 | 0.04 | −0.02 | 0.07 | 0.30 |
| **Local temperature** | −0.41 | 0.07 | −0.51 | −0.32 | 1.00 |
| Local precipitation | 0.00 | 0.04 | −0.03 | −0.04 | 0.27 |
| *Age at fledging* |  |  |  |  |  |
| **Year** | −0.03 | 0.02 | −0.05 | −0.01 | 0.76 |
| **Stopover freeze days** | 0.26 | 0.11 | 0.07 | 0.40 | 0.95 |
| Local precipitation | −0.05 | 0.09 | −0.18 | 0.01 | 0.39 |
| **Clutch initiation** | −0.18 | 0.12 | −0.31 | −0.02 | 0.75 |
| Brood size | 0.08 | 0.10 | −0.01 | 0.20 | 0.43 |

**Table A4.** Annual sample sizes for the clutch initiation date and age at fledging analysis. Due to low sample size, 2007 and 2010 were removed from consideration for age at fledging. Predicted daily nest survival (DNS) and survival estimates (± SE) are for first nests only. Survival is based on a 24-day nesting cycle (from first egg to fledge). The standard error was estimated by bootstrapping the predictions 1000 times.

| Year | Clutch initiation (n) | Age at fledging (n) | DNS | Survival (%) |
| --- | --- | --- | --- | --- |
| 2003 | 14 | 8 | $0.987 \pm 0.003$ | $72.8 \pm 3.7$ |
| 2004 | 15 | 13 | $0.997 \pm 0.002$ | $93.6 \pm 3.0$ |
| 2005 | 45 | 24 | $0.993 \pm 0.001$ | $84.1 \pm 1.8$ |
| 2006 | 55 | 19 | $0.978 \pm 0.001$ | $58.8 \pm 1.5$ |
| 2007 | 19 | 0 | $0.492 \pm 0.281$ | $00.0 \pm 2.5$ |
| 2010 | 21 | 1 | $0.898 \pm 0.002$ | $07.7 \pm 0.4$ |
| 2011 | 25 | 3 | $0.950 \pm 0.002$ | $29.4 \pm 1.4$ |
| 2015 | 56 | 13 | $0.952 \pm 0.001$ | $30.9 \pm 0.8$ |
| 2016 | 53 | 14 | $0.956 \pm 0.001$ | $34.4 \pm 0.9$ |
| 2017 | 35 | 10 | $0.968 \pm 0.001$ | $45.6 \pm 1.3$ |
| 2018 | 32 | 6 | $0.947 \pm 0.001$ | $27.3 \pm 0.7$ |
| 2019 | 12 | 7 | $0.991 \pm 0.002$ | $80.6 \pm 3.7$ |
| Total | 382 | 118 | $0.967 \pm 0.002$ | $44.2 \pm 1.2$ |

**Figure A1.**
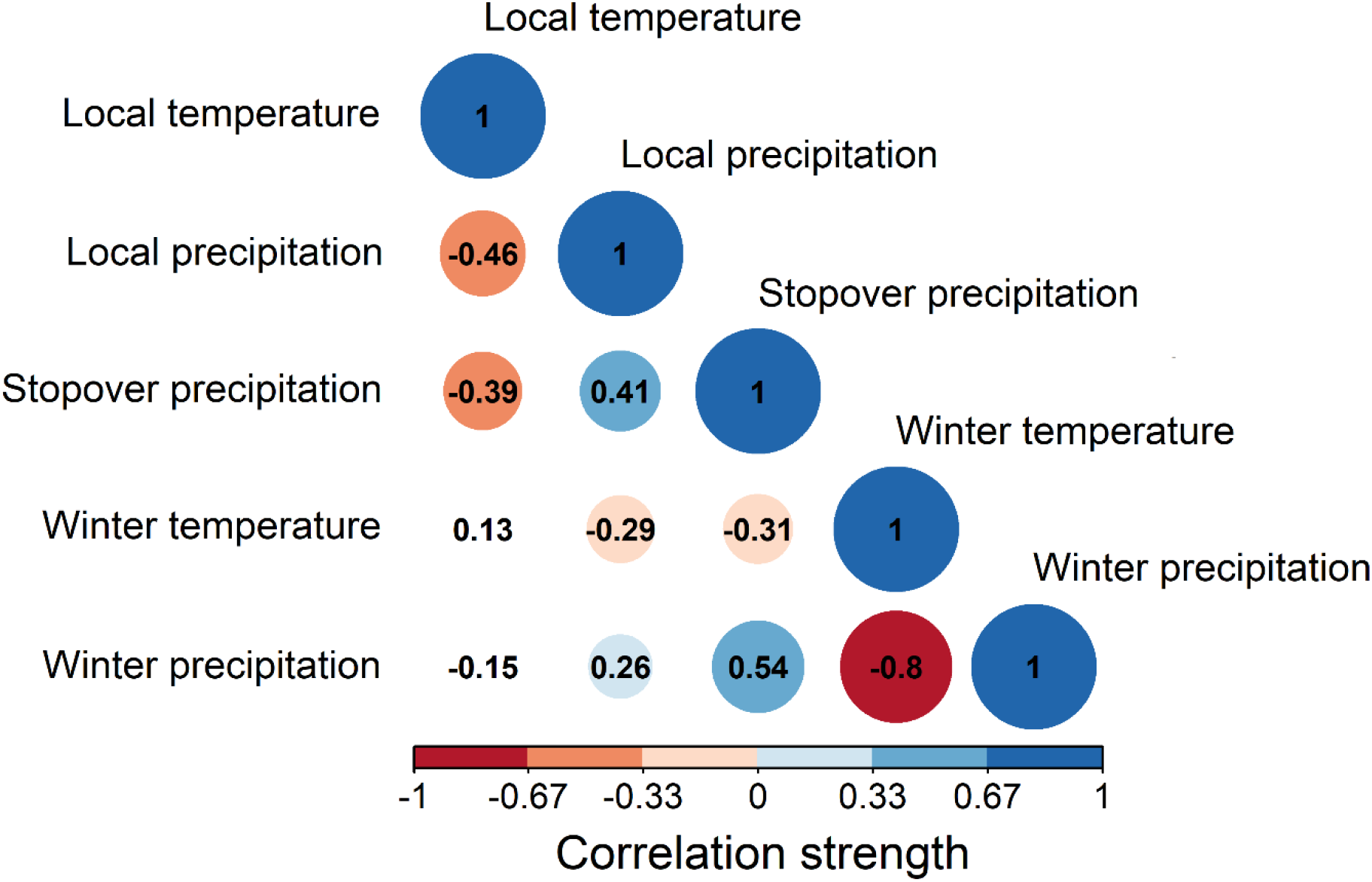
Pearson’s correlation matrix among the candidate climate variables for clutch initiation date. Larger, darker circles are stronger correlations, while the absence of a circle indicates weak to no correlation. A correlation greater than ±0.70 is considered a concern for collinearity.

**Figure A2.**
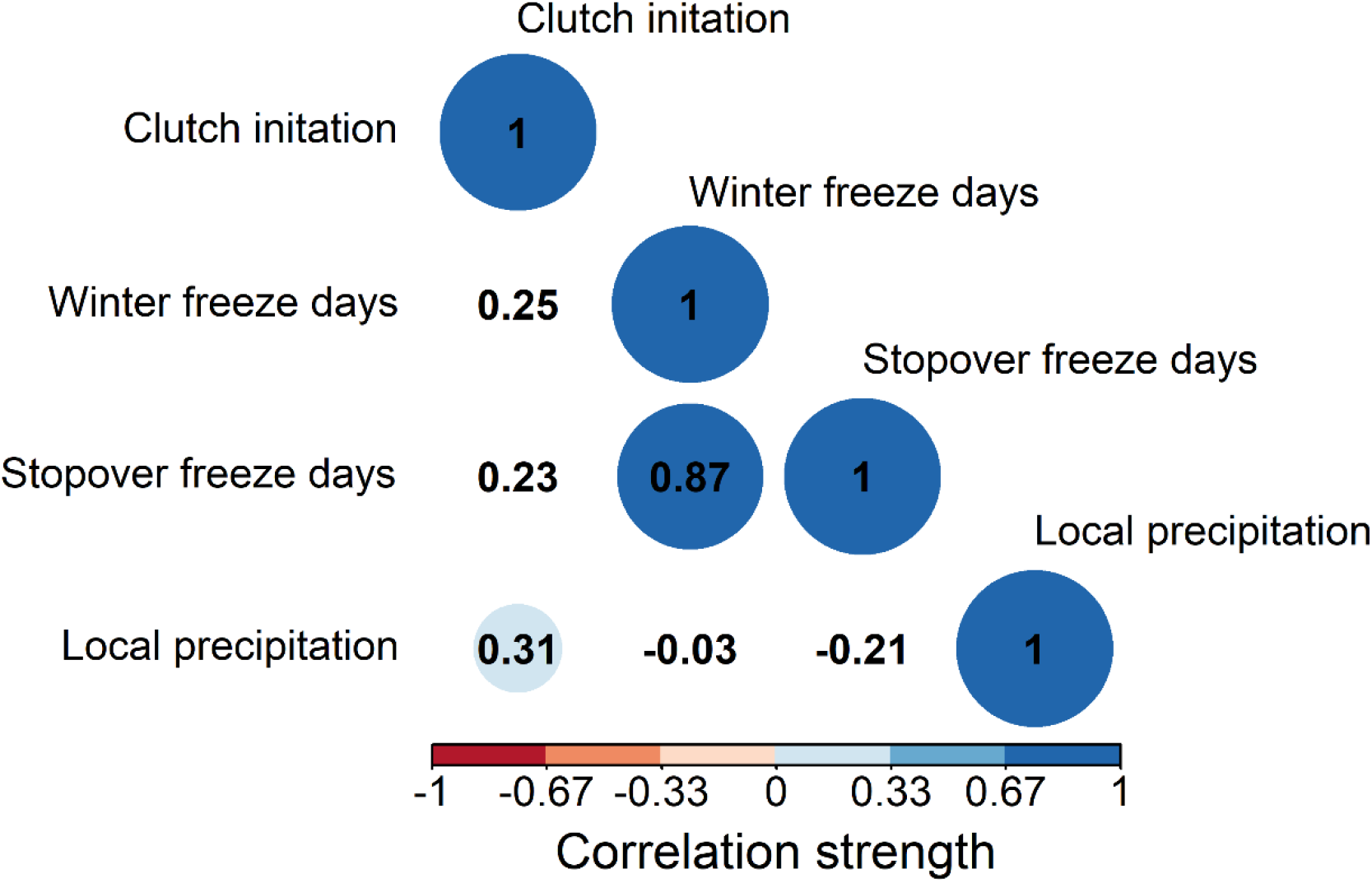
Pearson’s correlation matrix among the candidate climate variables for age at fledging. Clutch initiation was included to test for potential multi-collinearity with the climate variables. Larger, darker circles are stronger correlations, while the absence of a circle indicates weak to no correlation. A correlation greater than ±0.70 is considered a concern for collinearity.

## Notes

### Competing Interest Statement

The authors have declared no competing interest.

